# Thalamo-frontal connectivity patterns in Tourette Syndrome: Insights from combined intracranial DBS and EEG recordings

**DOI:** 10.1101/2024.10.09.617413

**Authors:** Laura Wehmeyer, Juan C. Baldermann, Alek Pogosyan, Fernando Rodriguez Plazas, Philipp Alexander Loehrer, Leonardo Bonetti, Sahar Yassine, Katharina Zur Mühlen, Thomas Schüller, Jens Kuhn, Veerle Visser-Vandewalle, Huiling Tan, Pablo Andrade

## Abstract

Thalamic deep brain stimulation (DBS) has shown clinical improvement for patients with treatment-refractory Tourette Syndrome (TS). Advancing DBS for TS requires identifying reliable electrophysiological markers. Recognising TS as a network disorder, we investigated thalamo-cortical oscillatory connectivity by combining local field potential (LFP) recordings from the DBS thalamic target region using the Percept^TM^ PC neurostimulator with high-density EEG in eight male TS patients (aged 27 to 38) while stimulation was off. We identified a spatially and spectrally distinct oscillatory network connecting the medial thalamus and frontal regions in the alpha band (8–12 Hz), with connectivity strength negatively correlated with TS symptom severity. Moreover, reduced thalamo-frontal alpha connectivity before tic onset, localised in sensorimotor regions and the inferior parietal cortex, suggests its direct role in tic generation. Importantly, associations with symptoms and pre-tic dynamics were specific to connectivity patterns and not evident in the pure power spectra. These findings underscore the importance of investigating electrophysiological oscillatory connectivity to characterise pathological network connections in TS, potentially guiding stimulation-based interventions and future research on closed-loop DBS for TS.

## Introduction

Tourette Syndrome (TS) is a neurodevelopmental disorder characterised by motor and vocal tics, often co-occurring with other neuropsychiatric conditions, including attention-deficit hyperactivity disorder (ADHD) and obsessive-compulsive disorder (OCD).^1^ Unlike other hyperkinetic disorders, tics are typically preceded by a premonitory urge (PMU) and can be voluntarily suppressed for a limited period.^2, 3^ It is widely accepted that dysregulations within the cortico-basal ganglia-thalamo-cortical (CBGTC) circuits contribute to the pathophysiology of TS.^4^ For patients with TS, deep brain stimulation (DBS) has emerged as a promising and safe treatment option. DBS of various targets within CBGTC circuits, particularly the thalamus, has demonstrated effectiveness in alleviating TS symptoms.^5–7^ However, the mechanism of DBS is still not fully understood, although, there is a growing consensus that DBS in general may exert its therapeutic effects by modulating network activity within CBGTC circuits.^8, 9^ To further advance and personalise stimulation-based treatment approaches for individuals with TS, it is crucial to develop a comprehensive understanding of the fundamental pathophysiological network mechanisms that should be the primary target of intervention.

Previous research has leveraged the unique opportunity offered by DBS to record intracranial local field potentials (LFPs) in patients with TS. These investigations have unveiled pathological low-frequency activity in the thalamus (range: 2–15Hz) associated with tic severity and tic generation, suggesting its potential as a biomarker for TS.^10–20^ While offering valuable insights, research on LFPs in TS is hampered by small sample sizes and predominantly conducted within intraoperative settings, introducing the surgery-induced microlesion effect and anaesthetics or analgesics as confounding factors.^21^ Furthermore, two studies have indicated that resting low-frequency power may normalise or simply change over time following DBS,^12, 15^ limiting the long-term applicability of these findings. Thus, there is a critical need for identifying a reliable and consistent biomarker that persists over the long-term after surgery.

Moreover, previous LFP research in TS primarily focuses on power characteristics and a notable gap persists in our understanding of the pathological connectivity between distant brain regions in patients with TS.^10–20^ Recognising TS as a network disorder, addressing this issue is crucial. Indeed, findings from neuroimaging,^22, 23^ stereotactic lesions,^24^ transcranial magnetic stimulation (TMS) research,^25^ and animal studies^26^ have highlighted the important role of CBGTC circuit connectivity in the TS pathophysiology. A powerful approach to characterise functional connections within CBGTC circuits is to assess electrophysiological oscillatory synchronisation between cortical and subcortical regions. This could be achieved by combining intracranial LFP with scalp recordings, such as EEG. However, since most prior LFP studies were confined to the intraoperative setting, the integration of scalp recordings was complicated due to open wounds and sterility.^27^

Addressing these limitations in prior LFP research in TS, advanced implanted neurostimulators with brain sensing capabilities, like the Percept^TM^ PC by Medtronic, now enable the recording of LFPs any time beyond the intraoperative phase.^28, 29^ This recently established, groundbreaking capability offers several advantages for research. Not only does it mitigate potential confounding effects from microlesions and enable the assessment of more naturalistic neural activity already subjected to DBS for an extended period, but it also greatly simplifies the integration of LFP with scalp recordings.

To date, no studies have been conducted using the Percept^TM^ PC in patients with TS to record LFPs, either exclusively or in combination with high-density EEG. Our study aims to address this gap by investigating thalamo-cortical oscillatory connectivity patterns in TS patients with implanted thalamic DBS systems, while stimulation was turned off. The principal correlate of functional connectivity investigated in this study is phase synchronisation, referring to the synchronisation of oscillatory phases between different brain regions. Our primary objective is to characterise oscillatory connectivity patterns at rest in terms of spatiality and spectrality and their association with TS symptom severity. Additionally, we intend to assess the impact of voluntary tic suppression on these spatially and spectrally segregated oscillatory connectivity patterns, and to investigate their potential dynamical changes in relation to the tic. These new insights may help to increase our understanding of electrophysiological markers related to TS symptoms and inform future research on closed-loop DBS for TS.

## Methods

### Participants

Eight adult patients with TS who underwent bilateral implantation of DBS electrodes in the medial thalamus, either the centromedian nucleus–nucleus ventrooralis internus (CM-Voi) or ventral anterior/ventral lateral nuclei (VA/VL), at the University Hospital Cologne between 2009 and 2022 were included in the present study. A detailed description of the surgical procedure targeting the CM-Voi in our centre can be found in Baldermann *et al.*^6^ and for the VA/VL target in Huys *et al*.^30^. Patients were implanted with either quadripolar or directional DBS leads from Medtronic, Minneapolis, USA. All patients received the Medtronic Percept^TM^ PC implantable pulse generator (IPG) either after DBS lead implantation or when the IPG had to be replaced due to battery depletion. Patients were clinically assessed at the time of testing using the Yale Global Tic Severity Scale (YGTSS)^31^ and Premonitory Urge for Tics Scale (PUTS-R)^32^. Please note that out of the initial eight patients included in the study, two had to be excluded from analysis due to excessive noise caused by frequent and intense tics, or excessive drowsiness and frequent eye closure during the experiment (Patient 4 & 5 - see Table 1 for individual characteristics of the final patient group). Each patient provided oral and written informed consent. The study was approved by the Ethics Committee of the Medical Faculty of the University of Cologne (No. 21–1351), registered in the German Clinical Trials Register (DRKS00029073), and performed in accordance with the Declaration of Helsinki.

**Table 1.** Patient Characteristics.

|  | Age | Sex | Months since OP | DBS Target | IPG Side | Coordinates | YGTSS |  | PUTS |
| --- | --- | --- | --- | --- | --- | --- | --- | --- | --- |
|  |  |  |  |  |  |  | Total | Global |  |
| Patient 1 | 27 | M | 26 | CM/Voi | Right | L: X: -5,00 Y: -4,00 Z: 0,00<br>R: X: 5,00 Y: -4,00 Z: 0,00 | 34 | 54 | 49 |
| Patient 2 | 33 | M | 85 | CM/Voi | Right | L: X: -5,00 Y: -4,00 Z: 0,00<br>R: X: 5,00 Y: -4,00 Z: 0,00 | 14 | 14 | 30 |
| Patient 3 | 27 | M | 3 | CM/Voi | Left | L: X: -5,01 Y: -4,00 Z: 0,00<br>R: X: 5,00 Y: -4,00 Z: 0,00 | 25 | 45 | 41 |
| Patient 6 | 24 | M | 3 | CM/Voi | Left | L: X: -6,00 Y: -4,00 Z: 0,00<br>R: X: 6,00 Y: -4,00 Z: 0,00 | 20 | 40 | 43 |
| Patient 7 | 38 | M | 164 | VA/VL | Left | L: X: -8,00 Y: -6,00 Z: -2,00<br>R: X: 8,00 Y: -6,00 Z: -2,00 | 21 | 41 | 35 |
| Patient 8 | 35 | M | 104 | CM/Voi | Left | L: X: -5,00 Y: -2,00 Z: 0,00<br>R: X: 5,00 Y: -2,00 Z: 0,00 | 11 | 36 | 22 |

Abbreviations: CM/Voi = Centromedian Nucleus–Nucleus Ventrooralis Internus; VA/VL = Ventral Anterior/Ventral Lateral Nuclei; IPG = Implantable Pulse Generator; L = Left; R = Right; YGTSS = Yale Global Tic Severity Scale; PUTS: = Premonitory Urge for Tics Scale.

### Experimental design

Recordings took place at 3 to 164 months after surgery 64.17 ± 64.69 (SD), with DBS turned off and medication unchanged. During the experiment, patients were seated comfortably in an armchair. Data was recorded at rest and during the Real-Time Urge Monitor task.^33^ At rest, patients were instructed to relax and to keep their eyes open or closed in alternating order for 7-9 minutes. Only data collected during the eyes-open condition were used for analysis. Patients were asked to avoid voluntary control over tics and express their tics freely during the rest recording. The Real-Time Urge Monitor task consisted of six 5-minute blocks with two alternating conditions: a ‘free-tic’ condition, similar to the rest recording, and a ‘tic-suppression’ condition, where patients were instructed to make their best effort to voluntarily suppress their tics. Moreover, patients were asked to indicate their PMU intensity in real time by moving a mouse on a scale from 0 to 100% displayed on the screen. The PMU ratings are reserved for future analysis as part of a separate investigation, and the present study focuses exclusively on the rest and tic-related data in line with our present study objectives. Of note, in patient 6, no rest recording could be performed, and only two blocks of the free-tic and no tic-suppression condition, because tics occurred only very infrequently during the recording, making tic suppression superfluous.

### Data recordings

EEG was recorded from 63 Ag/AgCl (EASYCAP GmbH, Herrsching, Germany) electrodes according to the extended 10–20 system using the ActiChamp amplifier and the BrainVision Recorder software (version 1.23.0003; Brain Products GmbH, Gilching, Germany). EEG data were online referenced to Cz, and a common ground was employed at FPz. Recordings were performed with a sampling rate of 5000 Hz and all impedances were kept below 10 kΩ. Additionally, EOGs (vertical and horizontal) and four accelerometers (1D acceleration sensors, Brain Products GmbH, Gilching, Germany) attached to the body parts of the most frequent tics were recorded using the same amplifier and recording software. A video was recorded, which was synchronised with the EEG using the Brain Vision Video Recorder software (Brain Products GmbH, Gilching, Germany). Simultaneously, bilateral LFPs were recorded with the Percept^TM^ PC using the programmer tablet connected to the communicator placed near the IPG.

Specifically, bipolar recordings for each hemisphere were performed in the BrainSense Streaming mode selecting the two outer contacts (0 and 3 for 1×4 and SenSight leads) as sensing contacts adjacent to the two middle contacts selected as stimulation contacts (1 and 2 for 1×4 leads; 1[a,b,c] and 2[a,b,c] for SenSight leads). LFPs were recorded while stimulation was turned off after a wash-out period of at least 2 minutes. Raw LFPs sampled at 250 Hz were low pass filtered at 100 Hz and high pass filtered at 1 Hz.^34^ To avoid long recording sessions that could potentially lead to data loss or export failures, a new streaming was started for the rest recording and each single block.^29^ During the streaming, the real time LFP was closely monitored for the presence of excessive artifacts and gaps in recordings. For later offline synchronization of EEG and LFP signals, DBS artifacts were induced in both LFP and EEG by briefly turning the stimulation on and off at the beginning and end of each streaming. Finally, the Percept’s JSON files containing the raw LFP data were exported from the programmer tablet for offline signal processing.

### Selection and marking of tic events

The video recordings were manually inspected offline for motor and vocal tics by an experienced clinician or psychologist using the VLC media player with millisecond precision (VideoLan, Paris, France). The start and end of each detected tic were marked in the EEG time series using Spike2 (Cambridge Electronic Design, Cambridge, UK). To ensure temporal precision, the timing of tics was cross verified using data from EOGs, accelerometers, and EEG. Subsequently, only tics preceded by a tic-free interval of at least 2 seconds were selected for analysis. Tics occurring in rapid succession with less than 2 seconds between them were considered part of a single tic sequence. In such cases, the start of the first tic in the sequence was marked as the start, and the end of the last tic as the end of the sequence. Because some patients exhibited a relatively low number of tics, tics recorded at rest and during the free-tic condition of the Real-Time Urge Monitor task were combined into a single free-tic condition. Consequently, the mean number of recorded tics per patient in the free-tic condition was 34.50 ± 32.13 (SD), with a range of 10–94 tics, and in the tic-suppression condition 32.20 ± 31.67 (SD), with a range of 11–87 tics.

### Signal processing

Data was pre-processed and analysed offline using custom-written Matlab scripts (Matlab 2023b, The Mathworks, Natick, MA, USA), Spike2 (Cambridge Electronic Design, Cambridge, UK), EEGLAB 2023.1^35^, and Fieldtrip (version 20230118; https://www.fieldtriptoolbox.org/). EEG data were bandpass filtered (fourth-order Butterworth IIR filter) from 1 to 50 Hz^36^ and resampled to 250 Hz to match the LFP sampling rate. The EEGLAB clean_rawdata function was employed to identify flatline, noisy, or outlier EEG channels, with no channels requiring removal. EEG data were then re-referenced to an average reference, and the Cz reference channel was reconstructed using spherical spline interpolation.^37^ Raw LFP data were extracted from JSON files using the Percept Toolbox, provided by Thenaisie *et al.* ^29^, with automatised correction for potential missing data points (available at https://github.com/YohannThenaisie/PerceptToolbox.git). However, LFP data from one tic-suppression block for patient 8 could not be extracted due to transmission problems. The extracted LFP recordings were checked for electrocardiogram (ECG) contamination and, if necessary (n = 1), cleaned of detected ECG artifacts through QRS interpolation using the Perceive Toolbox^38^ (available at https://github.com/neuromodulation/perceive). Subsequently, EEG and LFP signals were synchronised for each block/rest recording by manually aligning them with reference to the induced DBS artifacts or termination of high-frequency DBS, if visible in the signal, at the start of the recording.^29^ The synchronisation error range was determined by comparing the time difference between the synchronised EEG and LFP signals at the time of the second DBS artifact at the end of the recording. Due to an error range of 260 ms and 1000 ms, one free-tic block for patient 6, and another tic-suppression block for patient 8 had to be excluded from the analysis. For the remaining recordings, the mean error range was 7.29 ± 6.19 ms (SD) (range: 0 - 32 ms). Next, 5-second epochs were created around each tic onset, containing a 0.5 second buffer at both ends to avoid edge effects after spectral decomposition. Similarly, arbitrary non-overlapping 5-second rest epochs were extracted from all recordings by manually selecting time periods during which no tics or other movements, including mouse movements, occurred, with a minimum 2-second interval from any tic onset/offset. These rest epochs represent two distinct conditions: when derived from the free-tic condition, they reflect pure rest epochs, whereas epochs from the tic-suppression condition represent a state of tic suppression. For rest epochs, an automated artifact rejection based on extreme values was applied, considering only the time window of interest from −2 to 2 seconds around the arbitrary event ^39^. Epochs were rejected if EEG channel amplitudes reached a threshold of ± 200 μV. For tic epochs, each epoch was carefully manually inspected for artifacts in the time window before tic onset. Due to limited numbers of trials, patient 8 had to be excluded from the suppression analysis (n trials = 13) and patient 6 from the tic-related analysis (n trials = 8). The final trial numbers for each condition, along with the specification of which patient was included in which analysis, are provided in Table 2. Moreover, an extended infomax independent component analysis (ICA) was run on the segmented data. Independent components labelled as non-brain activity with at least 50% probability according to the IClabel algorithm^40^ were subsequently removed after visual inspection of the topographies of the first five components. This resulted in a total of 28.00 ± 8.49 (SD) retained independent components. Next, for each channel, epoched time series were decomposed from 2 to 40 Hz with a frequency resolution of 1 Hz (linear-spaced) using a second order IIR peaking resonator digital filter with a bandwidth of 2 at a 3 dB level, followed by Hilbert transformation. Subsequently, power and phase information were extracted for each epoch from −2 to 2 seconds to cut off the edge effects. Power was normalised by dividing through the mean power from 3 to 40 Hz and averaged in the theta (3-7 Hz), alpha (8-12 Hz), and beta (13-30 Hz) frequency bands.

**Table 2.** Trial Numbers and Analysis Inclusion Overview.

|  | Final Trial numbers |  |  |  |  | Included in ... |  |  |
| --- | --- | --- | --- | --- | --- | --- | --- | --- |
|  | Tic |  |  | Rest |  | Rest Analysis | Supp. Analysis | Tic Analysis |
|  | Free | Supp | All | Free | Supp |  |  |  |
| Patient 1 | 23 | 11 | 34 | 57 | 48 | Y | Y | Y |
| Patient 2 | 41 | 29 | 70 | 38 | 53 | Y | Y | Y |
| Patient 3 | 91 | 85 | 176 | 51 | 29 | Y | Y | Y |
| Patient 6 | 8 | ND | 8 | 30 | ND | Y | N | N |
| Patient 7 | 11 | 16 | 27 | 109 | 72 | Y | Y | Y |
| Patient 8 | 15 | 5 | 20 | 89 | 13 | Y | N | Y |

Abbreviations: Free = Free-Tic Condition; Supp= Tic-Suppression Condition; ND = Not Done; Y = Yes; N = No.

### Phase synchronisation

To quantify the functional connectivity between LFP and EEG signals across time, we calculated the phase synchronisation index (PSI) for each possible channel combination between the left and right thalamus and each single EEG channel. Specifically, the phase difference between signals was computed at all time points for each trial. The PSI over time was then calculated as the vector length of these phase differences within each trial (from −2 to 2 seconds) and subsequently averaged across trials. This yielded an index ranging from 0 to 1, with 0 indicating no synchronisation and 1 indicating perfect synchronisation. Similar to power, the PSI was then averaged in the theta, alpha, and beta frequency bands. Importantly, for the rest and suppression analysis, the PSI was calculated by computing the vector length of phase differences across the entire duration of epochs. Additionally, to determine the significance of phase synchronisation, the PSI of rest epochs solely derived from the free-tic condition was compared to surrogate phase-shuffled data. The surrogate data were generated 10 times using the same approach as described above, but with the order of LFP trials randomly shuffled, thereby destroying any phase synchronisation, and subsequently averaged.^27^ For the tic-related analysis, in order to investigate how cross region connectivity changes over time, the PSI was calculated within a sliding time window of 0.3 seconds moving in steps of 0.004 seconds (equal to one data point) from the start to end within each trial (from −2 to 2 seconds), to preserve the temporal dynamics associated with tics.

### Source Reconstruction

After preprocessing the EEG data, cortical source signals were reconstructed by solving the inverse problem, using FreeSurfer (available at http://surfer.nmr.mgh.harvard.edu), Brainstorm (available at http://neuroimage.usc.edu/brainstorm), and custom-written Matlab scripts. First, a template MRI (FSAverage) was used to obtain a 3D cortical mesh harboring 15,000 vertices.^41^ Subsequently, the Human Connectome Project (HCP) atlas was mapped to the cortical mesh, and a downsampled parcellation consisting of 26 regions covering each hemisphere was constructed.^42^ This process involved merging the 360 cortical areas from the HCP atlas into broader regions, while maintaining fine resolution within regions of the “somatosensory and motor cortex” and “paracentral lobular and mid cingulate cortex,” which are particularly relevant to TS based on existing literature.^2, 23, 24, 43–45^ Following this, EEG-channels and the template MRI were co-registered by matching individual digitised electrode positions with the surface data from the template MRI. The 3D positions of each EEG sensor on the scalp surface were digitised using an infrared dot-projection 3D scanner, specifically the Structure sensor Mark II (Occipital Inc., San Francisco, CA) integrated with an Apple iPad (8. Gen.).^46^ A realistic head model was then constructed using the OpenMEEG toolbox.^47, 48^ Finally, a linearly constrained minimum variance beamformer (LCMV)^49^ was employed to compute the regional time series data for the 52 cortical regions of interest from the epoched EEG data. In preparation for later tic-related analysis, power extraction and PSI calculation were conducted for cortical sources using the same methodology as for the EEG channels.

### Statistical analysis

Statistical analyses were performed at group-level on trial-averaged data using custom-written Matlab scripts. In our study, each thalamus was treated as one independent sample. Comparative analyses between conditions were performed using paired t*-*tests (Matlab function t-test) or multi-way analyses of variance (ANOVA) when testing fixed effects of multiple factors (Matlab function anovan). Significant ANOVA effects were followed by post-hoc pairwise comparisons (Matlab function multcompare). The normal distribution of the data was assessed using Shapiro-Wilk tests (Matlab function swtest). Since most of the data were not normally distributed, statistical significance for all statistical tests was determined by non-parametric Monte Carlo permutation tests.^50^ Specifically, p-values were derived by comparing the observed test statistic (i.e., F- or t-statistics) or difference between estimates in the case of pairwise comparisons with a distribution of statistics generated from shuffled data, created by randomly permuting the condition affiliation in 10000 permutation iterations. The p-value is then calculated as the proportion of shuffled test values that exceed the original test value, with the threshold for significance being established based on the top 5% of the distribution of permuted statistics. We applied correction for multiple comparisons using the false discovery rate (FDR) at α = .05 (Matlab function fdr_bh) or, for time series testing, cluster-based Monte Carlo simulations (MCS, α = .05, MCS p-value = .001).^51^ For the MCS analysis, p-values over time were binarized according to a threshold α = .05, and clusters of continuous significant values were identified. Binarized values were then shuffled within 10000 permutation iterations, generating a reference distribution of maximum cluster sizes for each permutation run. Original cluster sizes were finally compared against this reference distribution, with clusters exceeding the 99.9th percentile being considered significant.^51^ Beyond that, to examine relationships between variables, correlative analyses (Spearman’s correlations) or linear regression analysis were performed. All data are shown as mean ± SEM, unless otherwise indicated.

## Results

### A TS-protective Thalamo-Frontal Alpha Network at Rest

To characterise oscillatory connectivity patterns at rest, we conducted a rest analysis involving 6 patients (12 hemispheres), using arbitrary non-overlapping 4-second rest epochs derived from the free-tic condition. The normalised power spectrum of the averaged LFP across all hemispheres showed a gradual decrease with increasing frequency, culminating in a visually prominent low-frequency peak (Figure 1E). In the EEG, dominant alpha oscillations (8-12 Hz) most pronounced over posterior channels were visually observed (Figure 1G). To determine significant spatially and spectrally distinct thalamo-cortical phase synchronisation patterns, we compared the resting PSI between the LFP and each single EEG channel with surrogate phase-shuffled PSI data across different frequency ranges. For this purpose, we employed a three-way 2×3×2 ANOVA for each EEG channel, including Condition (original vs surrogate) and Frequency (theta vs alpha vs beta) as within-subject factors, and Hemisphere (left vs right thalamus) as a between-subject factor to account for potential lateralization effects. We were particularly interested in the interaction effect between Condition and Frequency, as this would indicate that differences in PSI between original and surrogate data vary across frequency bands. Such variation would highlight that certain frequencies show distinct phase synchronization patterns compared to others, indicating spectral specificity. FDR was applied across all p-values obtained for the single EEG channels and pairwise comparisons in case of post-hoc pairwise comparisons. Importantly, an interaction effect between Frequency and Condition was observed for frontal channels. Post-hoc pairwise comparisons revealed a significant difference between original and surrogate PSI for the depicted frontal channels only within the alpha frequency range (8-12 Hz), but not theta or beta (Figure 1A,B). This finding demonstrates a spatially and spectrally distinct thalamo-cortical phase synchronisation pattern specific to the alpha frequency band in frontal regions. Notably, no significant effects involving Hemisphere were observed, suggesting no lateralization effects, and justifying the treatment of left and right thalamus as independent samples.

**Figure 1:**
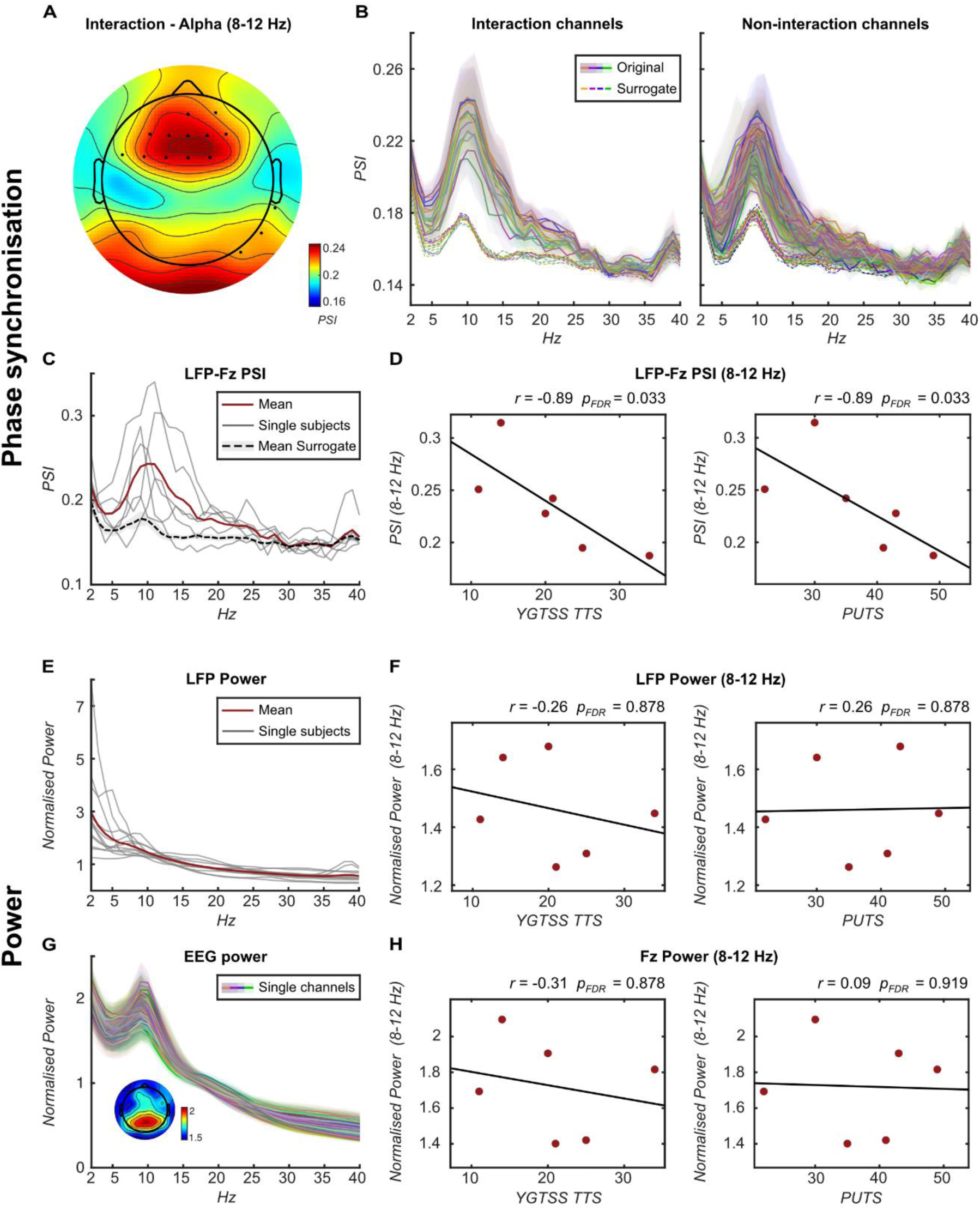
Resting thalamo-cortical connectivity and power patterns. (**A**) Topographic representation of the PSI calculated over 4-second rest epochs derived from the free-tic condition between the thalamus and single EEG channels (averaged across subjects) within the alpha frequency range (8-12 Hz). Dots indicate channels with an ANOVA interaction effect, where significant differences between original and surrogate PSI were observed only within the alpha, but not theta or beta frequency range. (**B**) Frequency plots illustrating original (solid line) and surrogate (dotted line) PSI between the thalamus and single EEG channels (averaged across subjects). Channels with significant interaction effects (corresponding to dots in panel A) are shown on the left, while other channels are displayed on the right. Shading represents standard error. (**C**) Frequency plot displaying the PSI between the thalamus and Fz, illustrating the original PSI for individual subjects averaged across hemispheres (grey lines), averaged across subjects (red line), and the surrogate PSI averaged across subjects (dotted black line). Shading represents standard error. (**D**) Scatterplots demonstrating Spearman’s correlations between PSI of thalamus and Fz (averaged across left and right thalamus) within the alpha frequency range and the YGTSS TTS (left) and PUTS (right). (**E**) Frequency plot showing normalised thalamic LFP power averaged over 4-second rest epochs derived from the free-tic condition for the individual subjects (grey lines) and the subject’s average (red line). Shading represents standard error. (**F**) Scatterplots illustrating Spearman’s correlations between normalised thalamic LFP power (averaged across left and right thalamus) within the alpha frequency range and the YGTSS TTS (left) and PUTS (right). (**G**) Frequency plot presenting normalised EEG power averaged over 4-second rest epochs derived from the free-tic condition for each channel averaged across subjects with a topographic representation of normalised power for single EEG channels within the alpha frequency range. Shading represents standard error. (**H**) Scatterplots illustrating Spearman’s correlations between normalised Fz power within the alpha frequency range and the YGTSS TTS (left) and PUTS (right). Abbreviations: YGTSS TTS = Yale Global Tic Severity Scale total tic score; PUTS = Premonitory Urge for Tics Scale.

Building upon this, we identified the maximum PSI within the alpha frequency range at Fz (PSI = 0.24 ± 0.02; Figure 1C), which was also significantly higher compared to the average PSI of all other channels within the same frequency range (*n* = 12, *t*_(11)_ = 3.26, *p* = 0.027, FDR-corrected). Consequently, we designated Fz and alpha as EEG channel and frequency range of interest for subsequent analyses. Next, to explore potential relationships with TS symptoms, we computed Spearman’s correlations between the alpha LFP-Fz PSI averaged across hemispheres and clinical parameters reflecting tic and urge severity (YGTSS total tic score (YGTSS TTS) and PUTS score, respectively), collected at the time of testing. We found a negative correlation between the alpha LFP-Fz PSI and tic/urge severity (*n* = 6; YGTSS TTS: Spearman’s Rho = −0.89, p = 0.033; PUTS: Spearman’s Rho = −0.89, p = 0.033; FDR-corrected; Figure 1D), indicating that higher phase synchronisation at rest was associated with less severe symptoms in our patient group. In contrast, no correlations were observed solely for thalamic and frontal alpha power (Figure 1F,H).

### Absence of a Tic Suppression Effect

To assess the impact of voluntary tic suppression, we performed a tic-suppression analysis involving 4 patients (8 hemispheres), comparing the free-tic condition with the tic-suppression condition. First, we compared tic frequency between conditions, excluding tics during rest recordings for comparability. No significant difference was found (*n* = 4, *t*_(3)_ = 0.16, *p* = 0.999; Supplementary Figure 1A). Similarly, when comparing arbitrary non-overlapping 4-second rest epochs derived from the free-tic condition with those from the tic-suppression condition, we found no effect on either the identified thalamo-frontal alpha phase synchronisation pattern or thalamic/frontal alpha power (LFP-Fz PSI: *n* = 8, *t*_(7)_ = −0.90, *p* = 0.381; Supplementary Figure 1B; LFP power: *n* = 8, *t*_(7)_ = −0.87, *p* = 0.412; Supplementary Figure 1C; Fz Power: *n* = 4, *t*_(3)_ = −1.01, *p* = 0.379: Supplementary Figure 1D). The absence of any tic suppression effect raises doubts about the effectiveness or presence of suppression.

### Tic-related Thalamo-Frontal Alpha Connectivity and Power Dynamics

To gain a comprehensive understanding of general tic-related neural patterns, tic epochs around tic onset from all conditions were pooled for analysis. Because no significant effects of tic suppression were observed and tic occurrence in each condition was limited, no further subgroup analysis was conducted for tics from the free-tic and tic-suppression conditions.

To capture dynamic changes in thalamo-frontal alpha phase synchronisation around tics, we conducted paired t-tests comparing LFP-Fz PSI values between tic and rest state within a 100-millisecond sliding time window, moving in steps of 20 milliseconds from −1.8 to 0.6 seconds relative to tic onset. Cluster-based multiple comparison correction revealed a significant PSI reduction from −0.22 to 0.18 s around tic onset (Figure 2A). To further explore the observed decreasing trend before tic onset, we performed a linear regression analysis to assess whether temporal changes in the PSI from −1.8 seconds to tic onset could be explained by time. The results indicated that time leading to the onset of tics accounted for 28% of the variation in PSI (*F*_(1,449)_ = 174.00, *p* < 0.001). Notably, employing the same cluster-based corrected sliding t-test approach to evaluate tic-related dynamics in thalamic/frontal alpha power, we observed a significant reduction in LFP power from −1.36 to −0.76 s before tic onset, but no significant changes in LFP or Fz alpha power immediately preceding the tic (Figure 2B,C). Furthermore, the variance in power before tic onset explained by time was relatively small in case of the LFP (*R²* = 0.11, *F*_(1,449)_ = 58.72, *p* < 0.001) or zero in the case of Fz (*R²* = −0.00, *F*_(1,449)_ = 0.00, *p* = 0.970). While these findings indicate a direct relationship between temporal changes in thalamo-frontal alpha phase synchronisation and tic generation before tic onset, they also suggest that thalamic/frontal alpha power by itself may not play a direct role in tic generation.

**Figure 2:**
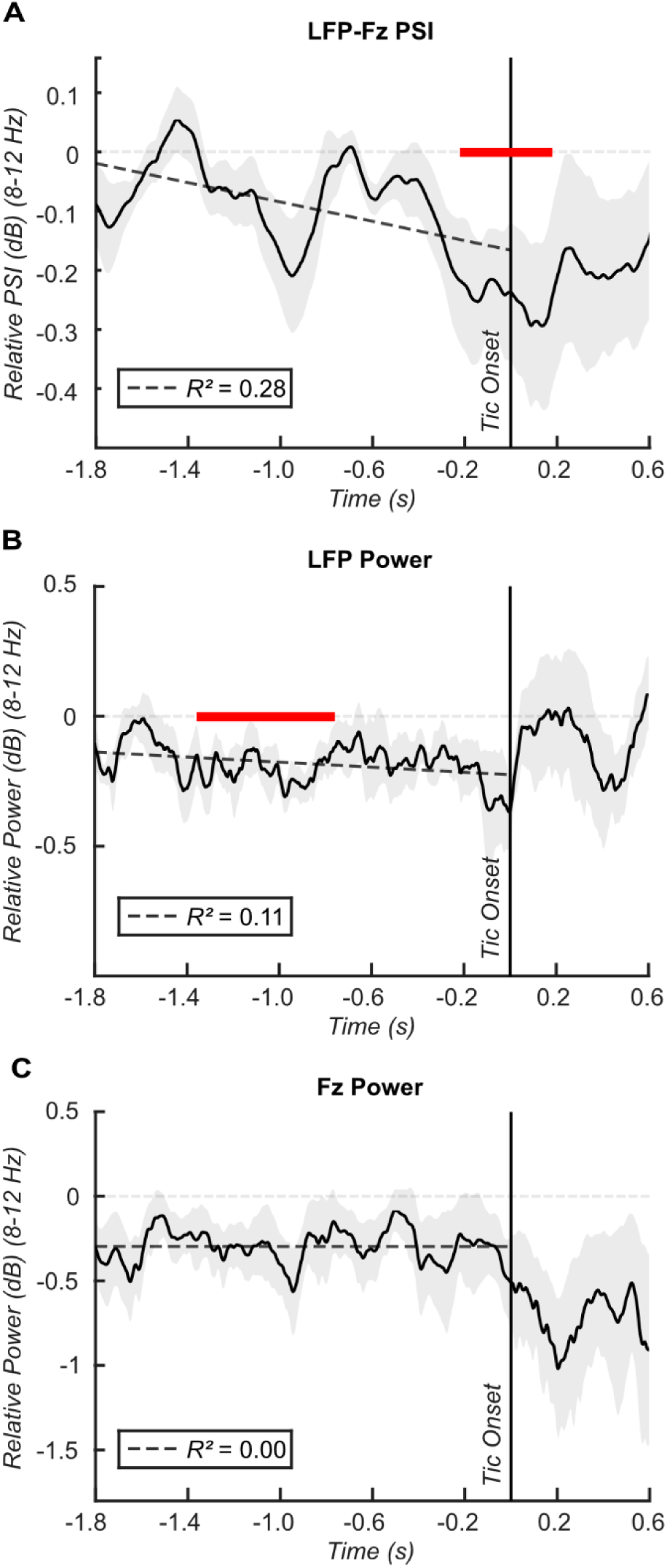
Tic-related thalamo-cortical connectivity and power dynamics at the sensor level. (**A**) Line plot illustrating the relative change in PSI from rest to tic between the thalamus and Fz within the alpha frequency range (8-12 Hz) averaged across subjects. PSI was calculated within a sliding time window of 0.3 seconds moving in steps of 0.004 seconds. Respective PSI values were compared between rest and tic state within a sliding time window of 0.1 seconds moving in steps of 0.02 seconds from −1.8 to 0.6 seconds relative to tic onset. Red bars indicate time windows of significant difference between tic and rest after cluster-based multiple comparison correction. A regression line from −1.8 to tic onset depicts the relationship between the relative PSI and time, along with corresponding R^2^ values in the lower left box. Shading represents standard error. (**B**) The corresponding line plot for relative change in thalamic LFP power. (**C**) The corresponding line plot for relative change in Fz power.

Expanding our investigation to the source-level, we aimed to better understand the spatial distribution of tic-related temporal changes in thalamo-frontal alpha phase synchronisation. We selected the following eight cortical regions of interest based on observed frontal modulation patterns at the sensor-level and their known relevance to tic generation in TS^2, 23, 24, 43–45^: Primary Motor Cortex (M1), Primary Somatosensory Cortex (S1), Cingulate Motor Cortex (CMC), Supplementary Motor Area (SMA), Premotor Cortex (PMC), Insular and Frontal Opercular Cortex (IC/FOp), Anterior Cingulate and Medial Prefrontal Cortex (ACC/mPFC), and Inferior Parietal Cortex (IPC). Using the same cluster-based corrected sliding t-test approach as described above, we uncovered varied patterns of tic-related temporal dynamics in phase synchronisation between the thalamus and these cortical sources (Figure 3). Specifically, connectivity between thalamus and S1, as well as IPC, showed a brief significant decrease starting around 1.3 seconds before tic onset (Figure 3E,G). This was followed by a broader reduction in connectivity involving the SMA, CMC, M1, PMC, IC/FOp, lasting until approximately 700 ms before tic onset (Figure 3B-D,F,I). Shortly after, a distinct decrease in connectivity to the ACC/mPFC was observed around 600 ms before tic onset (Figure 3H), followed by another short-lasting connectivity reduction to the SMA around 250 ms before tic onset (Figure 3B). Finally, connectivity between the thalamus and M1, S1, PMC, and IPC began to decrease again around 200 ms before tic onset, persisting until up to 180 ms after tic onset (Figure 3D-G). Additionally, regression analyses showed a gradual decrease in connectivity between the thalamus and these regions leading up to tic onset, with up to 62% of the variation in thalamus-S1 PSI changes accounted for by time (*F*_(1,449)_ = 722.16, *p* < 0.001; Figure 3E). Notably, no significant tic-related source power changes were observed (Figure 4).

**Figure 3:**
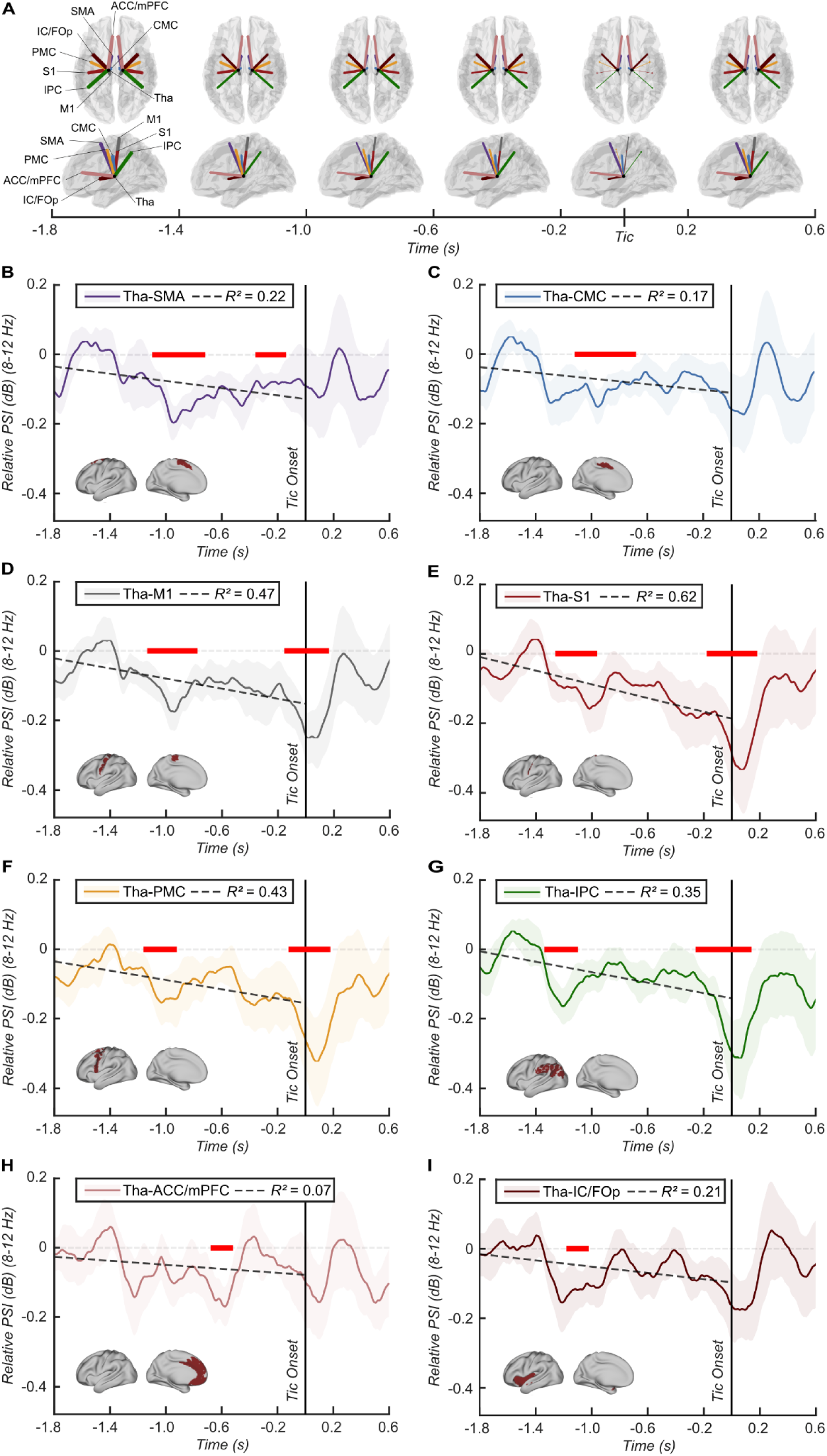
Tic-related thalamo-cortical connectivity dynamics at the source level. (**A**) Brain templates illustrating the relative change in PSI from rest to tic calculated using the sliding time window approach between the thalamus and eight selected sources of interest within the alpha frequency range (8-12 Hz), averaged across subject, and aggregated into 400 ms time windows from −1.8 to 0.6 seconds around tic onset. Colored lines depict connections between the thalamus and corresponding sources (illustrated for ipsilateral connections in both hemispheres for simplicity), with the thickness of each line representing the strength of the relative PSI in relation to the minimum and maximum PSI values across all sources and time windows. (**B-I**): Line plots showing the time series of the relative PSIs for the respective single sources. PSI values were compared between rest and tic state within a sliding time window of 0.1 seconds moving in steps of 0.02 seconds from −1.8 to 0.6 seconds relative to tic onset. Red bars indicate time windows of significant difference between tic and rest after cluster-based multiple comparison correction. A regression line from −1.8 to tic onset depicts the relationship between the relative PSI and time, along with corresponding R^2^ values in the upper box. Shading represents standard error. Brain templates in the lower left corners illustrate the spatial extent of the selected source. Abbreviations: SMA = Supplementary Motor Area; CMC = Cingulate Motor Cortex; M1 = Primary Motor Cortex; S1 = Primary Somatosensory Cortex; PMC = Premotor Cortex; IPC = Inferior Parietal Cortex; IC/FOp = Insular and Frontal Opercular Cortex; ACC/mPFC = Anterior Cingulate and Medial Prefrontal Cortex.

**Figure 4:**
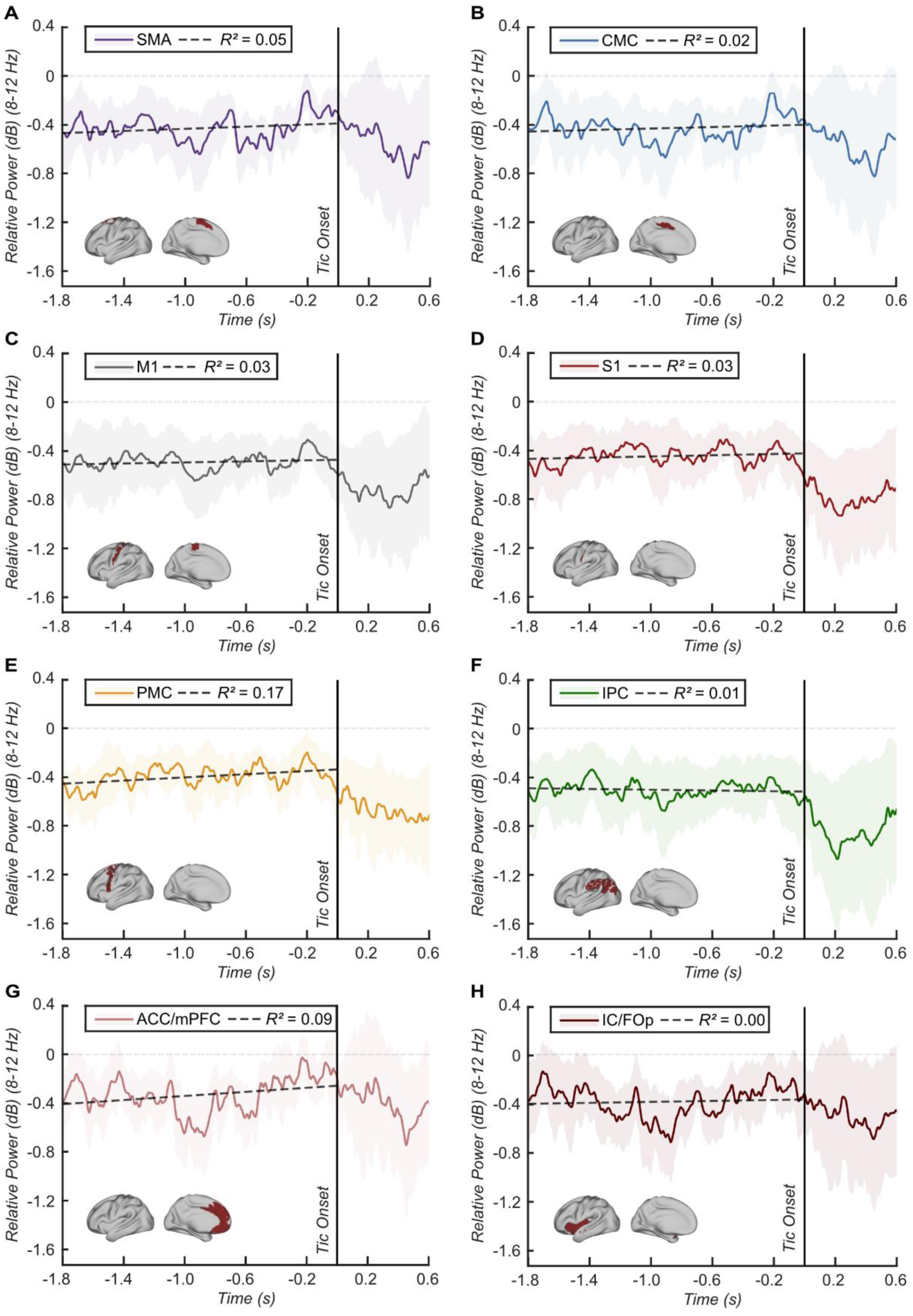
Tic-related thalamo-cortical power dynamics at the source level. (**A-H**): Line plots showing the time series of the relative change in power from rest to tic for each selected source of interest within the alpha frequency range (8-12 Hz) averaged across subjects. Power values were compared between rest and tic state within a sliding time window of 0.1 seconds moving in steps of 0.02 seconds from −1.8 to 0.6 seconds relative to tic onset. Red bars indicate time windows of significant difference between tic and rest after cluster-based multiple comparison correction. A regression line from −1.8 to tic onset depicts the relationship between the relative power and time, along with corresponding R^2^ values in the upper box. Shading represents standard error. Brain templates in the lower left corners illustrate the spatial extent of the selected source. Abbreviations: SMA = Supplementary Motor Area; CMC = Cingulate Motor Cortex; M1 = Primary Motor Cortex; S1 = Primary Somatosensory Cortex; PMC = Premotor Cortex; IPC = Inferior Parietal Cortex; IC/FOp = Insular and Frontal Opercular Cortex; ACC/mPFC = Anterior Cingulate and Medial Prefrontal Cortex.

## Discussion

This study represents the first endeavour to combine LFP recordings from the Percept^TM^ PC with high-density EEG, aiming to investigate thalamo-cortical oscillatory connectivity patterns via phase synchronisation in TS patients with an implanted DBS system in the medial thalamus. This innovative approach underscores the practicality of using sensing-enabled neurostimulators, exemplified by the Percept PC from Medtronic, for research purposes, enabling the acquisition of unique data otherwise unattainable, thereby offering invaluable contributions to our understanding of TS. Our findings revealed a spatially and spectrally distinct oscillatory network, connecting medial thalamus and frontal regions in the alpha (8– 12 Hz) band, with connectivity strength negatively correlated with TS symptom severity. Additionally, we demonstrated a reduction in thalamo-frontal alpha connectivity immediately preceding tic onset, suggesting its involvement in tic generation. Further analysis refining the spatiality of this finding revealed that this modulation extended to sensorimotor regions, including the primary motor cortex, primary somatosensory cortex, premotor cortex, as well as the inferior parietal cortex. Notably, pre-tic-related temporal dynamics are specific to phase synchronisation and not evident in the pure power spectra for both LFP and cortical sources.

Before proceeding to the discussion of our main findings, it is essential to address the absence of a tic suppression effect in the present patient group. While our findings indicate no significant impact on tic frequency from voluntary suppression, we lack additional data related to tic suppression efforts for further insights. Therefore, only speculations can be made about possible explanations. One possibility is that patients did not voluntarily suppress their tics when requested to do so, for various reasons. Some patients may find tic suppression too challenging because they cannot resist the urge.^52^ In other patients, symptom severity may have been sufficiently low to render tic suppression superfluous or impractical, given the difficulty in detecting impending tics due to low urge intensity.^53^ However, another possibility is that tics were inadvertently suppressed even in the free-tic condition. Individuals with TS often develop habitual tic control mechanisms, particularly in social settings, implying that tic suppression may occur automatically over time.^52, 54, 55^ Consequently, tic control may persist even when patients are not actively attempting to suppress their tics. Therefore, we cannot rule out the potential influence of (automatic) tic control in the supposed to be free-tic condition. Beyond that, it should be noted that the small number of patients (n=4) included in the suppression analysis likely challenged the detection of a tic suppression effect.

Unique to the present study is the comprehensive characterisation of thalamo-cortical connectivity patterns across different frequency ranges covering the entire cortex. We discovered a spatially and spectrally distinct thalamo-cortical network in patients with TS at rest, restricted to the alpha frequency band (8–12 Hz) in frontal regions. Notably, simultaneous resting cortical alpha power peaking in posterior, rather than frontal, regions suggests that the observed thalamo-frontal alpha connectivity pattern is independent of overall power activity. Interestingly, we observed a negative correlation between thalamo-frontal alpha connectivity and tic/urge severity. At the same time, no similar correlation pattern could be observed for thalamic and frontal alpha power, emphasising the distinctiveness of the relation between the identified connectivity pattern and TS symptomatology. This finding underscores the importance of considering TS as a network disorder characterised by pathophysiological functional connections within CBGTC circuits. It aligns with prior neuroimaging findings of abnormal connections between the thalamus and various frontal regions, encompassing motor and sensory cortices, the cingulate cortex, and the supplementary motor area.^22, 56^

The specific mechanism underlying the observed association between increased thalamo-frontal alpha connectivity and reduced symptom severity remains speculative. One plausible hypothesis is that increased thalamo-frontal connectivity could potentially enhance (automatic) tic control, which is supported by previous research linking fronto-striatal hyperconnectivity as well as general cortical alpha network connectivity to chronic tic control.^54, 57^ Also, it has been postulated that tic control involves top-down control mechanisms originating from frontal to subcortical regions, potentially normalising abnormal activity within CBGTC circuits responsible for tics.^52, 58, 59^ However, considering that thalamo-frontal alpha connectivity also negatively correlated with urge severity, an alternative or complementary hypothesis could be that increased connectivity may be associated with a reduced PMU. This would also be in line with the observed dynamical decrease of thalamo-frontal alpha connectivity preceding tic execution, as discussed later. Given the thalamus’ role in sensorimotor function as a central mediator of sensory input and perception it is reasonable to posit that thalamo-frontal connections may influence the PMU.^60, 61^ Moreover, previous research has highlighted the critical role of frontal regions in the PMU.^2, 62, 63^

Although the precise mechanisms are yet to be fully understood, the negative correlation between thalamo-frontal alpha connectivity and symptom severity suggests its potential as a target for stimulation-based treatments in patients with TS. Consistent with this, neuroimaging studies have shown that DBS is most effective when structural or functional connectivity networks linking the thalamus to the frontal cortex, particularly sensorimotor regions such as the (pre-)SMA, cingulate cortex, primary motor cortex, and primary sensory cortex, are stimulated.^64–67^ Furthermore, research utilising median nerve stimulation (MNS) highlights the importance of targeting the alpha frequency range, as rhythmic 10-Hz pulse trains have shown significant tic improvement.^68, 69^ Rhythmic 10-Hz median nerve stimulation may increase the thalamo-frontal alpha connectivity, which may reduce the occurrence of tics. Interestingly, we observed dynamic changes in thalamo-frontal alpha connectivity in relation to the tic. These were characterised by distinct connectivity decreases between the thalamus and frontal regions, particularly in sensorimotor areas and the inferior parietal cortex, at different timings before the tic. A notable reduction in connectivity around one second before the tic involved various brain regions, including the SMA, cingulate motor cortex, primary motor cortex, primary somatosensory cortex, premotor cortex, inferior parietal cortex as well as the insular and frontal opercular cortex. This indicates that neural processes underlying tic occurrence start well before tic onset, which is in line with the typically observed pre-tic symptomology in TS patients, i.e. the PMU.^2^ Furthermore, immediate connectivity decreases around tic onset involved the primary motor cortex, primary somatosensory cortex, premotor cortex, and inferior parietal cortex. This finding is particularly interesting as it implies a direct link to tic generation. Importantly, these immediate pre-tic changes were specific to connectivity patterns, with no similar direct tic-related dynamic changes detected for mere thalamic or frontal alpha power at either sensor or source level.

Our understanding of the precise mechanisms underlying these pre-tic disconnections remains speculative. Building on our earlier hypothesis regarding the nature of the resting thalamo-frontal connectivity pattern, the observed disconnection immediately preceding tics might indicate a transient lapse in tic control, potentially facilitating tic execution. However, the temporal pattern of gradually decreasing connectivity over time leading up to the tic suggests more a progressive development of underlying processes, culminating in the manifestation of the apparent tic. Such a process could be more likely related to the PMU, which typically increases before the tic until reaching its peak just before tic onset.^2^ It is also plausible that the decreases observed around one second before the tic and immediately before tic onset represent different underlying processes. In fact, the present connectivity patterns may stem from a complex interplay of processes involving both tic control and PMU, engaging different brain regions at different timings.

The observed tic-related dynamic connectivity changes, encompassing different sensorimotor, frontal, and parietal brain areas are in line with various observations from imaging studies on tic-preceding neural activity.^23, 43, 44^ Previous LFP studies have primarily focused on tic-related thalamic power changes, consistently reporting a distinct unrhythmic low-frequency (2-10 Hz) increase following tic onset.^10, 13, 14, 16, 18^ Based on this feature, closed-loop DBS approaches in TS have already demonstrated feasibility, safety, and efficacy comparable to continuous DBS.^19, 20^ However, these studies did not identify any pre-tic activity changes. Similarly, a recent EEG study found no pre-tic alterations in alpha or beta power in sensorimotor cortices, contrasting with the well-known movement-related beta suppression observed before voluntary movements.^70^ This highlights the absence of a distinct electrophysiological power marker preceding tic onset, suggesting the involvement of a complex neural network in tic generation. Prior electrophysiological research on tic-related thalamo-cortical connectivity patterns in TS is very limited.^10, 18^ One study combining chronic LFP recordings with surface electrocorticogram (ECoG) recordings over the motor cortex detected no thalamo-motor cortex coherence during rest, movement, or tics, which could be related to the limited coverage provided by subdural strips.^18^ In another study, intraoperative combined LFP and EEG recordings in three patients revealed repetitive increases in thalamo-cortical coherence preceding tics across broad frequency ranges, including alpha and beta.^10^ Discrepancies between these findings and ours may be attributed to factors such as the timing of the recordings and potential cross-subject variability.

In light of this, our results add valuable insights to the existing literature by demonstrating a consistent pattern of pre-tic-related connectivity changes across patients, extending beyond the intraoperative time window. These findings may pave the way for future research aimed at identifying electrophysiological pre-tic markers, particularly for closed-loop DBS in TS.

Various limitations of the present study need to be acknowledged. First, it was limited by a small and heterogeneous patient sample. In terms of gender, the sample demonstrated homogeneity with all participants being male. Next, correlation results may be influenced by DBS effects, as clinical parameters reflect symptom severity over the past week when DBS was active. To accurately assess the relationship between thalamic activity and symptom severity in the DBS-Off state, it would be necessary to collect clinical parameters after turning off DBS for at least a week. However, this is unfeasible due to clinical and ethical constraints. A further limitation may arise from potential synaptic plasticity changes following long-term stimulation, as effects of stimulation administered over months might not fully reverse within a wash-out period of 2 minutes. In our tic-related analysis, a major limitation is the lack of a control condition for comparison, such as voluntary movements. Additionally, we cannot rule out the potential influence of other movements during the pre-tic state, as patients preformed mouse movement as part of the task. The considerable heterogeneity in the phenomenological appearance of tics introduces another limitation. However, it should be emphasised that our primary aim was to identify a common neural substrate underlying tics, irrespective of their specific characteristics.

In conclusion, the present study, combining LFP recordings using the Percept^TM^ PC with high-density EEG in TS patients with thalamic DBS, extends beyond previous intraoperative LFP studies, providing valuable new insights. Our findings implicate the role of a distinct thalamo-frontal network within the alpha frequency band (8–12 Hz) in the TS pathophysiology. Thereby, they underscore the importance of investigating electrophysiological oscillatory synchronisation between subcortical and cortical regions to characterise pathological functional connections within CBGTC circuits. These identified connectivity patterns may serve as targets for stimulation-based interventions in TS, informing future research on closed-loop DBS for TS.

## Acknowledgements

We wish to thank the participating patients for making this study possible, Annika Sauter for helping with the recordings, and the rest of the MRC BNDU Tan group for providing useful discussions on data analysis.

## Data availability

The raw data are not yet openly available, due to data privacy regulations of patient data. We will consider requests to access the data in a trusted research environment as part of a collaboration. Please contact the corresponding authors for this.

