## Supplementary Material for "Thalamo-frontal connectivity patterns in Tourette Syndrome: Insights from combined intracranial DBS and EEG recordings"

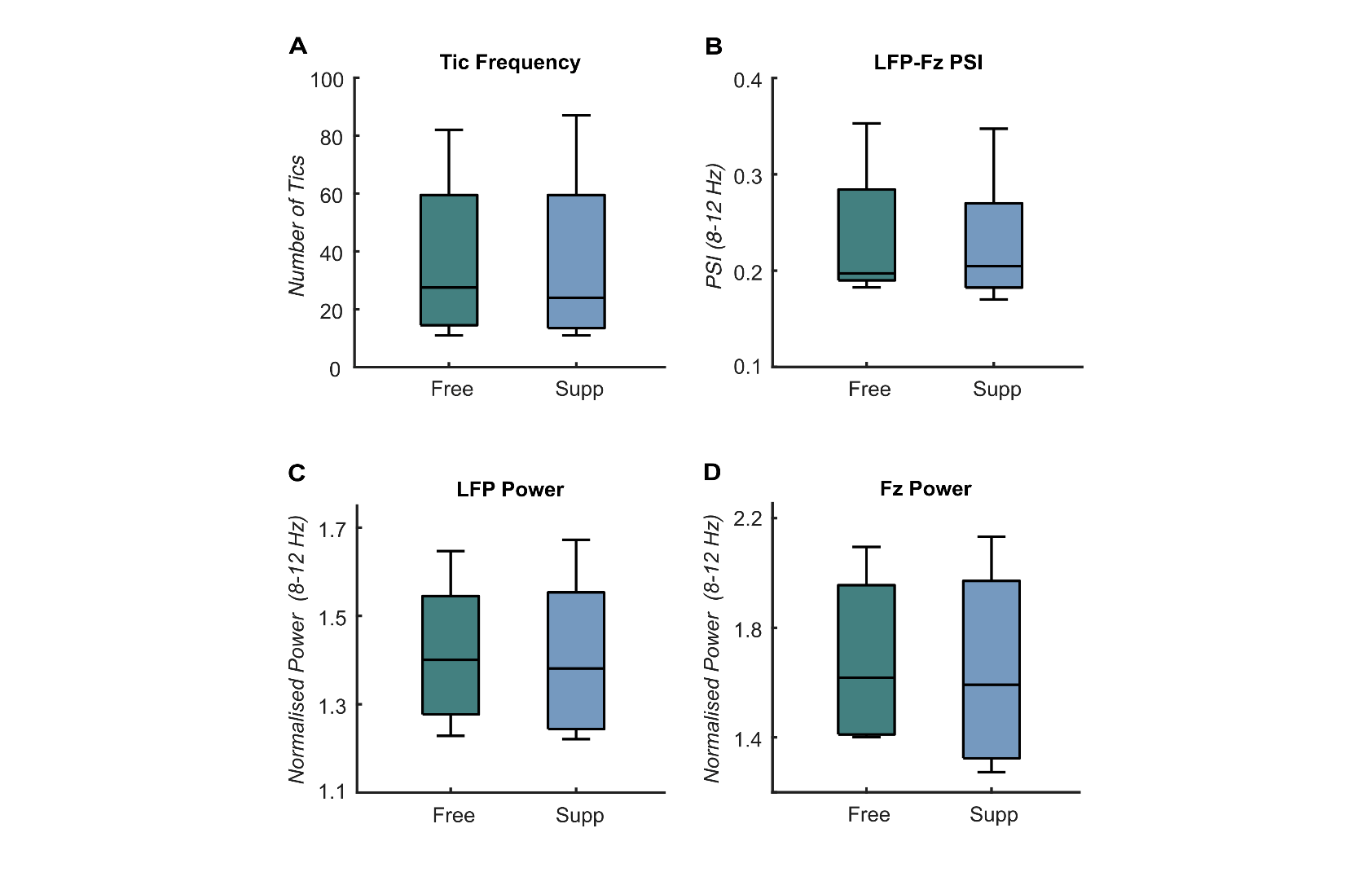

**Supplementary Figure 1: Effect of tic suppression.** Boxplots showing the group-level difference between free-tic and tic-Suppression conditions for (**A**) The number of tics; (**B**) The PSI calculated over 4-second rest epochs derived from the free-tic condition and tic-suppression condition between the thalamus and Fz within the alpha frequency range (8-12 Hz); (**C**) Normalised thalamic LFP power averaged over 4-second rest epochs derived from the free-tic condition and tic-suppression condition within the alpha frequency range; (**D**) Normalised Fz power averaged over 4-second rest epochs derived from the free-tic condition and tic-suppression condition within the alpha frequency range. Abbreviations: Free = Free-tic condition; Supp = Tic-suppression condition.
